# Atomic-level characterization of protein-protein association

**DOI:** 10.1101/303370

**Authors:** Albert C. Pan, Daniel Jacobson, Konstantin Yatsenko, Duluxan Sritharan, Thomas M. Weinreich, David E. Shaw

**Affiliations:** D. E. Shaw Research, New York, NY 10036, USA.; Department of Biochemistry and Molecular Biophysics, Columbia University, New York, NY 10032, USA.

**Author notes:** To whom correspondence should be addressed. Albert C. Pan, David E. Shaw.

## Abstract

Despite the biological importance of protein-protein complexes, determining their structures and association mechanisms remains an outstanding challenge. Here, we report the results of atomic-level simulations in which we observed five protein-protein pairs repeatedly associate to, and dissociate from, their experimentally determined native complexes using a new molecular dynamics (MD)-based sampling approach that does not make use of any prior structural information about the complexes. To study association mechanisms, we performed additional, conventional MD simulations, in which we observed numerous spontaneous association events. A shared feature of native association for these five structurally and functionally diverse protein systems was that if the proteins made contact far from the native interface, the native state was reached by dissociation and eventual re-association near the native interface, rather than by extensive interfacial exploration while the proteins remained in contact. At the transition state (the conformational ensemble from which association to the native complex and dissociation are equally likely), the protein-protein interfaces were still highly hydrated, and no more than 20% of native contacts had formed.

---

Most proteins associate with other proteins to function, forming complexes that lie at the heart of nearly all physiological processes, including signal transduction, DNA repair, enzyme inhibition, and the immune response. Determining the structures of these complexes and elucidating their association mechanisms are problems of fundamental importance. While substantial progress has been made toward the structural determination of protein-protein complexes, such structures are still relatively underrepresented in the protein data bank,^1^ especially compared to the large number of known, functional protein-protein interactions derived from high-throughput, non-structural approaches like yeast two-hybrid screening and affinity purification-mass spectrometry.^2^ Moreover, the structures of many complexes that are important drug targets for cancer and autoimmune disease remain difficult to determine experimentally.^3,4^ As for protein-protein association mechanisms, powerful experimental approaches like double-mutant cycles and paramagnetic relaxation enhancement (PRE) have afforded a wealth of information about potential transition states and intermediates,^5,6^ but these data are often indirect or limited to, for example, metalloproteins or proteins with attached paramagnetic spin labels. Obtaining direct, atomic-level detail about association pathways for a diverse set of protein-protein complexes and developing broader insights into the common principles of protein-protein association mechanisms are still open problems.

Atomic-level molecular dynamics (MD) simulations offer a computational route toward characterizing both the structure and dynamics of protein-protein complexes. Using MD, one could in principle start a simulation with two protein monomers and “watch” them associate and dissociate reversibly during a single trajectory. Such a simulation would provide an unprecedented sampling of the possible complexes that can be formed by the protein monomers and a straightforward way to rank the stability of different complexes based on that fraction of the time during which each complex is observed. Moreover, mechanistic details like intermediate and transition states along the association pathway could be observed. In practice, however, it has proven difficult to study protein-protein association and dissociation in MD simulations: Only a few examples have been reported^7–15^ of simulations that successfully captured spontaneous protein-protein association to an experimentally determined complex, and these examples have been limited to examining one system, or to the association of smaller peptides.^7,11,14^ Reversible association during a single simulation trajectory has not been observed at all.

Difficulties encountered in such simulations include the formation of non-native associated states with long lifetimes compared to simulation timescales. Such kinetic traps severely hamper the sampling of other states—including states that may be thermodynamically more favorable, and thus more likely to represent the most populated complex at physiological conditions.^16,17^ Even if the most thermodynamically stable complex is sampled, observing spontaneous dissociation could require simulations on the order of seconds to days^18^. Some approaches have attempted to overcome this timescale problem by combining data from multiple short simulation trajectories,^10,12,14^ and have had success in modeling various aspects of protein-protein association. None of these methods, however, has been applied to a broad set of structurally diverse protein-protein systems, making it difficult to draw general conclusions about protein association. Moreover, such methods require an additional layer of modeling to combine the short trajectories, and necessarily rest on assumptions that can bias the results toward atypical pathways or miss important conformational states.^19,20^

In this paper, we have used long-timescale MD simulations in combination with a newly developed enhanced sampling approach, which we call *tempered binding*, to simulate the reversible association of five structurally and functionally diverse protein-protein systems, and have also performed conventional MD simulations to capture many spontaneous association events. In a tempered binding simulation, the strength of interactions between the protein monomer atoms, and sometimes between the protein monomer and solvent atoms, is scaled at regular time intervals using a simulated Hamiltonian tempering framework^21–23^. This scaling allows long-lived states to dissociate more quickly. In practice, tempered binding involves a conventional MD simulation augmented by frequent Monte Carlo moves that update the scaling strength among rungs on a ladder of values. The Monte Carlo updates are detail-balanced such that, at each rung of the ladder, a Boltzmann distribution of states corresponding to that rung’s value of the scaling factor is properly sampled. In particular, the sampling at the lowest rung of the ladder (rung 0) corresponds to the completely unscaled Hamiltonian and is consistent with the distribution of states sampled in a conventional MD simulation. Further methodology details are available in the SI.

We applied tempered binding to six protein-protein systems and observed reversible association in five out of the six systems (Figs. S1 and S4; Table S1). For the five reversibly associating systems, the most stable complex in the simulation agrees with the complex determined crystallographically within atomic resolution (Fig. 1). (In the sixth protein-protein system, the protein dimer CLC-ec1,^24^ we observed association to the experimentally determined complex, but did not observe dissociation on the timescales of our simulations (Fig. S4).) Some of these simulations also sampled alternative bound states that could have functional relevance, and provided quantitative estimates of the free energy difference between the native state and these alternative states (Figs. 2, S2, and S3). Although for this initial study we have limited ourselves to proteins that do not undergo large conformational changes upon binding (Table S1), we note that such proteins in themselves constitute a large class of important protein-protein complexes^25^. (The ribonuclease HI-SSB-Ct system could be considered an exception, since the SSB-Ct peptide folds upon binding to the ribonuclease, but the RMSD difference between the folded and disordered forms of the peptide is only about 2 Å).

**Figure 1.**
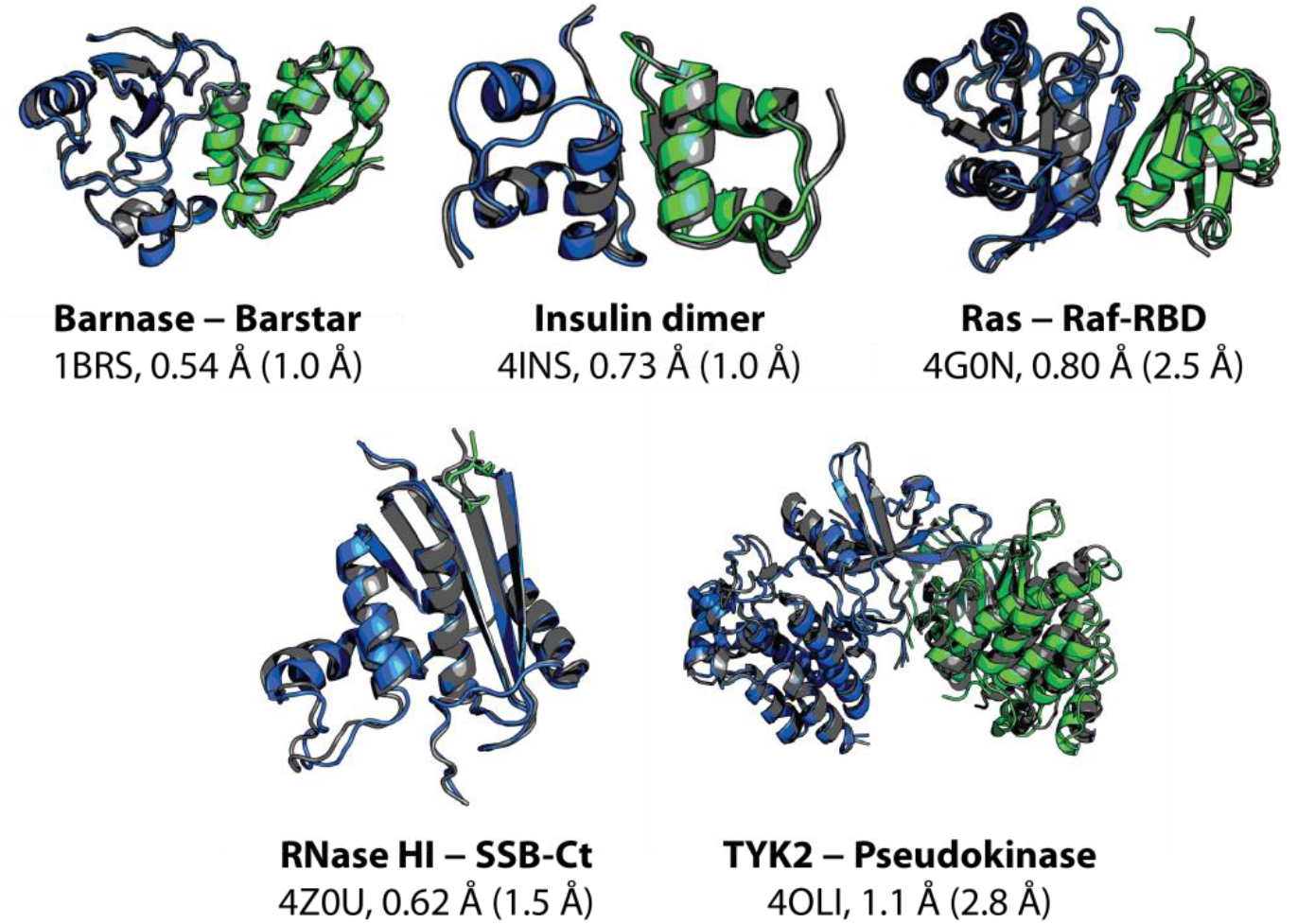
The most thermodynamically stable complex visited during reversible-association simulations agrees with the experimentally determined complex within atomic resolution. Representative structures of the most thermodynamically stable complexes observed in reversible-association simulations are shown. For each protein-protein complex, we show a representative associated structure obtained from simulation (blue and green) superimposed on the experimentally determined crystal structure (gray) by a best-fit Cα alignment of the larger protein monomer (blue), along with the name of the complex, the PDB entry of the experimental structure,^3,26,36–38^ and the Cα interface and ligand root-mean-square deviations (I-RMSD and L-RMSD) between the two structures. The protein-protein interface is defined as any pair of Cα atoms, one from each protein monomer, within 10 Å of each other. The L-RMSD is calculated by first aligning the Cα atoms of the larger protein monomer and then calculating the Cα RMSD of the smaller protein monomer (green).^1^ Tempered binding simulations for these five protein-protein systems used the Amber ff99SB*-ILDN^39–41^ force field and the TIP3P^42^ water model. The representative structure was obtained by clustering the simulations, to avoid bias toward the experimentally determined structure. Clustering was only performed on simulation frames sampled at the lowest tempering rung (rung 0), where the distribution of states is the same as that of a conventional MD simulation. The representative structure from the cluster with the greatest occupancy is used in the figure. Because the tempered binding approach scales all interactions uniformly, the simulations were not biased for, or steered toward, any particular protein-protein complex. Although tempered binding successfully recapitulated experimentally determined bound structures for these systems, its computational expense greatly exceeds that of other approaches, such as docking, for this particular task. We note, however, that its accuracy for this set of five complexes is considerably better than a current state-of-the-art docking program, particularly in its ability to select the correct native-like structure among various low-energy protein-protein complexes (Table S4), speaking both to the level of sampling achieved by tempered binding and to the accuracy of current MD force fields. Additional descriptions of the systems and the methods are available in the SI.

**Figure 2.**
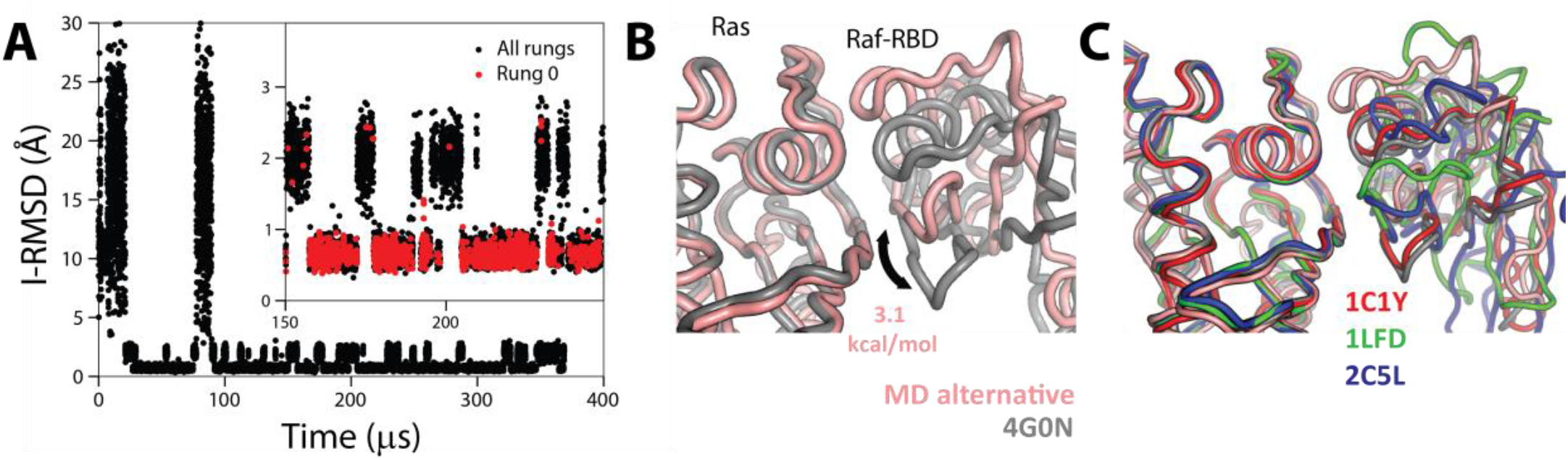
Tempered binding provides a direct atomic-level observation of the ensemble of bound states involved in protein-protein interactions. (A) An I-RMSD trace of a tempered binding simulation of the Ras protein binding to the Ras-binding domain of the Raf effector protein (Raf-RBD) shows reversible association to the crystal complex (PDB ID: 4G0N)^37^. For Ras-Raf-RBD, the simulation not only reached the known crystal-structure complex, which was the most thermodynamically stable state, but also an alternative complex about 2 Å away from the crystal structure. The inset shows a portion of the RMSD trace zoomed in along the y-axis. Black (red) circles are points from all (rung 0) trajectory frames. (B) A structure of the alternative state (pink) overlaid onto the crystal structure (gray), aligned to the Ras domain, is shown. Counting the population of the crystal-like state versus the alternative state in rung 0 suggests that the alternative state is approximately 3.1 kcal mol^−1^ higher in free energy than the crystal-like state. (C) An overlay of other Ras-effector complexes demonstrates that the conformation of this alternative state is well within the range of observed binding-interface conformations.^43–45^

Our current tempered binding protocol (which focuses on scaling the near electrostatic interactions between protein monomers, between protein monomers and water molecules, and, in the case of CLC-ec1, between protein monomers and lipid molecules), resulted in a significant increase in sampling efficiency in our simulations of the protein-protein systems studied in this work. In a tempered binding simulation of the enzyme-inhibitor system barnase-barstar, for example, the protein-protein system escaped from its native complex in 100s of μs (Fig. S1), whereas the lifetime of the native complex is on the order of a day,^26^ a speedup of almost nine orders of magnitude. It is possible, however, that other tempered binding protocols might further improve sampling efficiency and perhaps allow us to observe reversible association for the CLC-ec1 dimer (Fig. S4).

In addition to the tempered binding simulations, we performed hundreds of conventional MD simulations of the five reversibly associating protein systems in order to study their association mechanisms (Table S3). In each of these simulations, we observed the proteins come into contact and form loosely associated protein-protein configurations (encounter complexes) that then either (a) formed the specific, close-range interactions in the native complex without the proteins at any point dissociating (a successful association event), (b) dissociated without first reaching the native complex (an unsuccessful association event), or (c) remained kinetically trapped in a non-native state for the remainder of the simulation. (Here, we use the term “encounter complex” to refer to the set of protein-protein configurations in which any heavy atom in one protein is within 4.5 Å of any heavy atom in the other protein, but in which the interface RMSD is not within 1.5 Å of the native complex.) In some simulations, there were unsuccessful association events preceding a successful association event. We note that we observed several successful association events for each of the five systems (Table S3), and, as expected, we observed no examples of dissociation once the experimentally determined complex formed.

Successful association events in these five systems shared several key features. Rather than forming an encounter complex at a random interface and reaching the native interface (without dissociating) by way of an extensive search, in successful association events the encounter complexes tended to form near the native interface, at least for events observed within the timescale of our simulations (~10s of μs). (For a given protein-protein system, the contact preceding a successful association event tended to be nonspecific, varying among different events, but typically involved interactions between charged residues in or close to the native binding interface.) In contrast, encounter complexes that formed during *unsuccessful* association events displayed a wide variety of relative protein-protein positions, with no pronounced preference for their positions in the native complex (Figs. 3 and S5).

**Figure 3.**
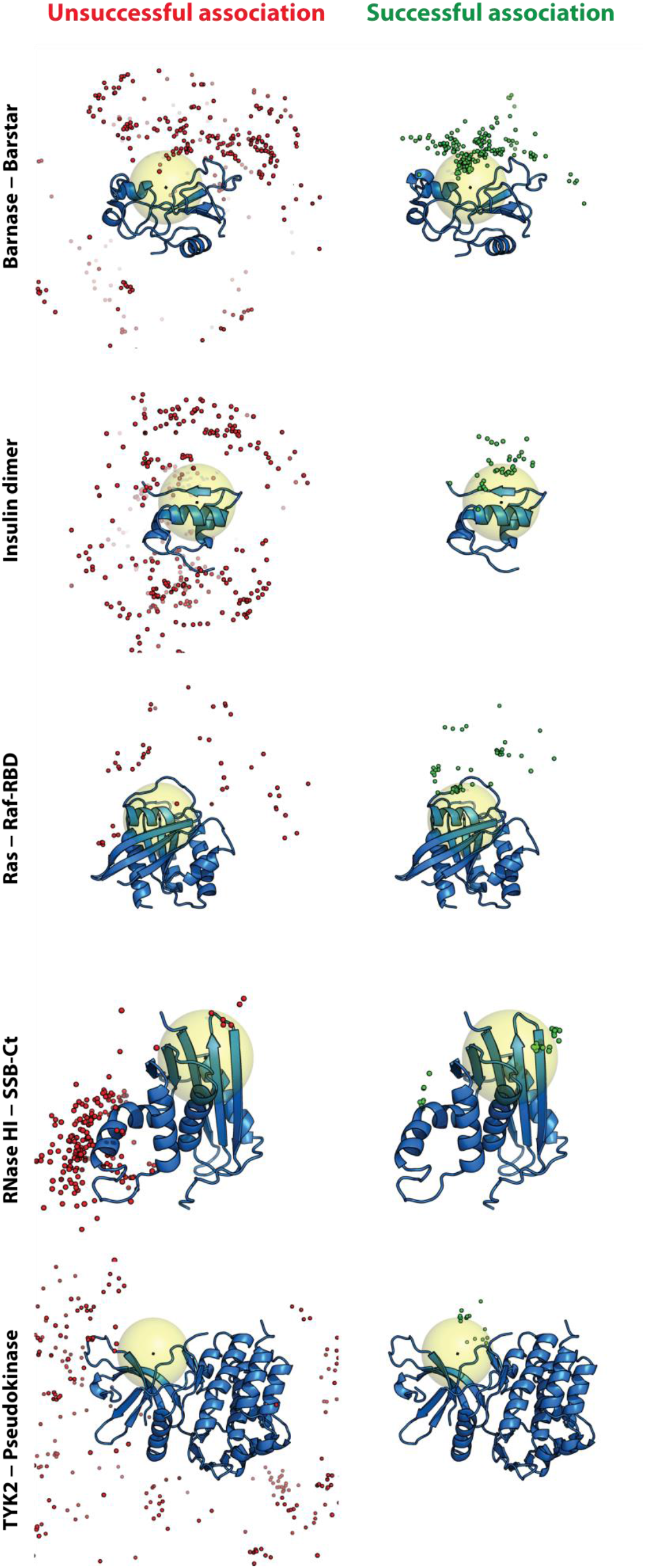
Encounter complexes visited in successful association events favored structures in which the two proteins were positioned similarly to how they are positioned in the experimentally determined complex. Simulation frames were uniformly sampled from encounter complexes and aligned to the larger protein. A single snapshot of the larger protein (blue cartoon) is shown for reference, overlaid with multiple snapshots of a Ca atom of the smaller protein near the center of the native binding interface taken from unsuccessful (red spheres) and successful (green spheres) association trajectories. The large yellow sphere indicates a region defined by a 10-Å radius around the center of mass of the binding interface of the larger protein. Kinetically trapped non-native states, which neither dissociated nor reached the native state during a simulation, were not included in this analysis.

Successful association events in the five systems also shared similar features at the transition state (the ensemble of configurations from which association and dissociation are equally likely): No more than 20% of native contacts had formed in these configurations, and the protein-protein interfaces were still highly hydrated. We identified configurations in the transition state ensemble of association by calculating the probability of successful association, *p_Assoc_* (also known as the committor probability), for several configurations during a successful association Event,^27–28^ and were able to identify configurations at or near the transition state for all five systems. A configuration is classified as a member of the transition state ensemble if *p_Assoc_* = 50% (that is, if additional trajectories initiated from that configuration with randomized velocities drawn from a Boltzmann distribution commit half of the time to the native complex and half of the time to the unbound state). All the transition states characterized here occurred when <20% of native contacts had formed and while there were still a significant number of water molecules between atoms that are in contact in the native complex (Fig. 4). Given the intensive computational effort required for determining committor probabilities and identifying transition states, we only determined committor values in one successful association event for each system condition. The general features of the transition state configurations were found to be qualitatively similar among simulations of barnase-barstar with different force fields (Table S3), and even among completely different systems (Fig. 4D), however, providing strong evidence that these transition states are representative of the transition state ensembles for these five systems.

**Figure 4.**
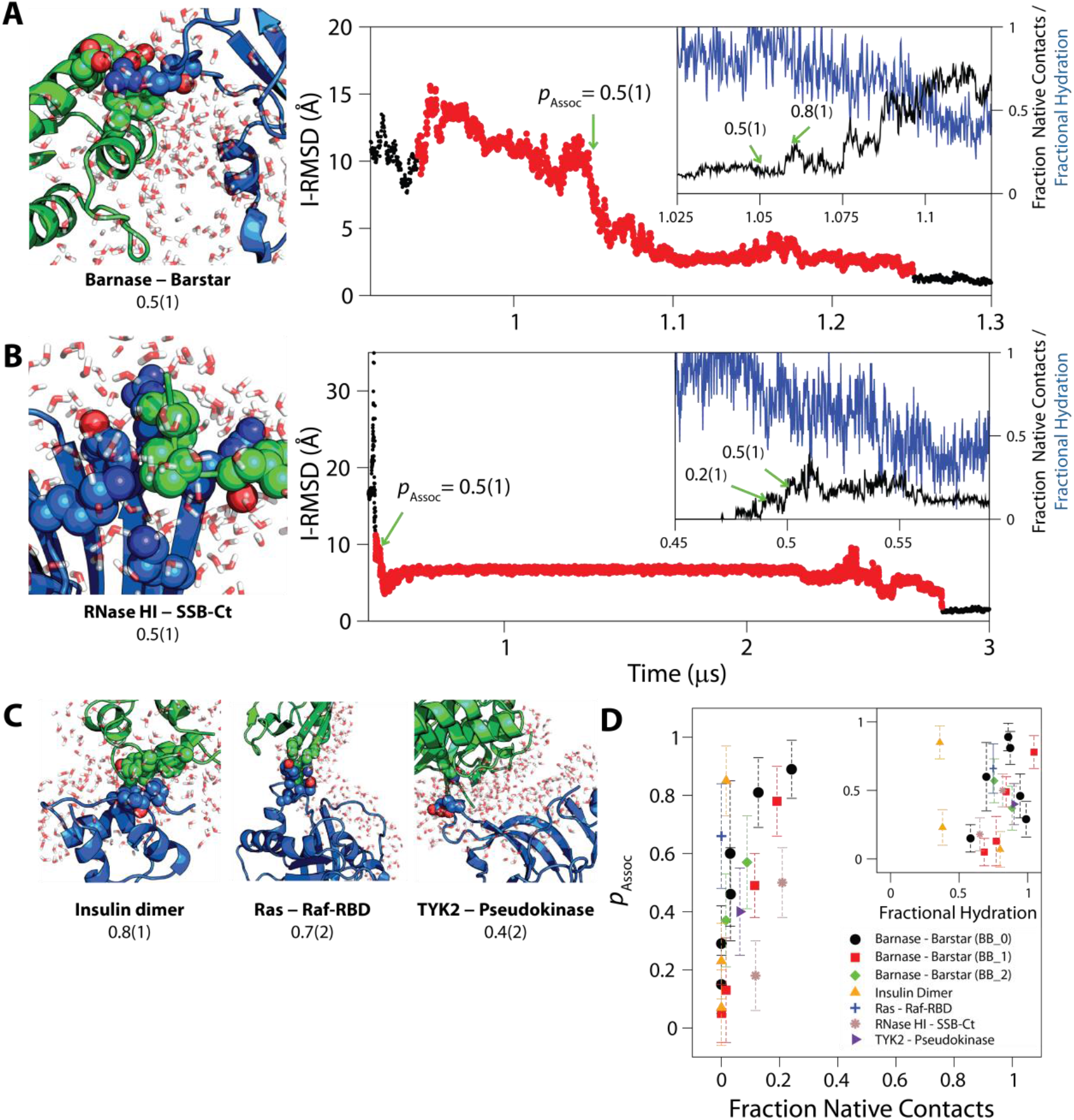
The transition state for association was solvated, and only <20% of native contacts had formed. Configurations and portions of I-RMSD traces from successful association events in conventional MD simulations of (A) barnase-barstar and (B) RNase HI-SSB-Ct association are shown. The green arrows indicate points where p_Assoc_ was calculated in the successful association event (red circles). The insets show an expanded view of the region close to the transition state, showing the fraction of native contacts (black) and fractional hydration around the interface (blue) as a function of time (see SI). Additional configurations near the transition state of association are shown for (C) insulin dimer, Ras-Raf-RBD, and TYK2-pseudokinase. Configurations show the larger protein as a blue cartoon, the smaller protein as a green cartoon, inter-protein residue contacts between residues at the native interface as van der Waals spheres, and water within 4 Å of the native binding interfaces as red and white licorice, demonstrating the lack of protein-protein contacts and the large amount of water at the binding interface. (D) A plot of the probability of association, *p_Assoc_*, against the fraction of native contacts and a normalized water interface coordinate (inset). The fraction of native contacts remained below 30% even for configurations that had a committor probability of 90% to the native-complex state. Protein-protein interfaces in the successful association events, except for insulin dimer, which has a relatively hydrophobic interface, also remained more than 50% solvated.

For the enzyme-inhibitor complex barnase-barstar, one of the best experimentally characterized protein-protein complexes,^26^ our simulations are consistent with previous experimental and computational work, including extensive mutational analysis^5,29^ and Brownian dynamics simulations.^30,31^ Combining the information from the tempered binding and conventional MD simulations, we estimated the binding free energy, Δ*G*_b_, to be 19.2(2) kcal mol^−1^ (*K*_D_ = 1.0(2) × 10^−14^ M), the association rate, *k*_on_, to be 2.3(2) × 10^7^ M^−1^s^−1^, and the dissociation rate, *k*_off_, to be 2.3(3) × 10^−7^ s^−1^. These simulation-derived values correspond relatively well with the known experimental values of Δ*G*_b_ = 19 kcal mol^−1^, *k*_on_ = 6 × 10^8^ M^−1^ s^−1^, and *k*_off_ = 8 × 10^−6^ s^−1^.^32^ (Further discussion of how the simulation values were calculated can be found in the SI.)

Notably, our atomic picture of the transition state agrees with mutational and kinetic studies from Schreiber, Fersht, and co-workers, which suggest that the transition state occurs before a majority of native interactions are formed, and while the protein-protein interface is still highly Solvated.^5,33^ In addition, the observation that associating proteins already had relative positions similar to those in the native complex upon making contact during successful simulated association events is consistent with the idea that long-range electrostatic attraction is involved in the rapid association of barnase-barstar^31,34^ and other protein-protein pairs with oppositely charged binding sites^35^. The strong affinity of barstar for barnase and the hydrophilic nature of the interface make it a relatively unusual protein-protein system, so it is striking that the features of the transition state and encounter complex observed in our simulations were common to the association mechanisms of all of the protein-protein systems studied in this work (Fig. 4). This shared mechanism may thus also apply to the broader class of protein-protein complexes that— like the systems studied here (Table S1)—do not undergo large conformational changes upon binding.

We have observed reversible association of a set of five protein-protein systems to their respective experimentally determined structures using a new enhanced sampling method that enabled an increase in sampling efficiency of as much as nine orders of magnitude. Together with our long-timescale conventional MD simulations, which yielded many spontaneous association events, our results provide an atomic-level view of protein-protein association mechanisms. In the future, this methodology could be used to determine the structures and association mechanisms of at least some protein-protein complexes that have not yet been experimentally characterized. The ability to observe both association and dissociation events could be especially useful in this context, helping to distinguish thermodynamically stable complexes from kinetically trapped states that are sparsely populated.

## Acknowledgments

The authors thank Michael Eastwood, Cristian Predescu, Jesus Izaguirre, and Doug Ierardi for helpful discussions, and Berkman Frank, Rebecca Bish-Cornelissen, and Jessica McGillen for editorial assistance.

